# HEXOKINASE-dependent regulation of WRKY transcription factors in Arabidopsis

**DOI:** 10.1101/2023.01.29.526134

**Authors:** Joshua M. Boyte, Runyu Xie, Yandong Liu, Xiang Li, Christopher R. Buckley, Michael J. Haydon

**Affiliations:** School of BioSciences, University of Melbourne, Parkville, VIC 3010, Australia

**Keywords:** Sugar signalling, superoxide, reactive oxygen species, metabolism, HXK1

## Abstract

Sugars are the major product of photosynthesis and provide the stored energy and basic building blocks for all living cells. Sugars also act as dynamic signals throughout the plant life cycle to regulate growth, development and interactions with the biotic and abiotic environment. From a previous RNA-seq experiment, we have identified eight sugar-regulated *WRKY* transcription factor genes. Focusing on four, we find that *WRKY11, WRKY17, WRKY60* and *WRKY72* are upregulated by sucrose, glucose or fructose by a superoxide signalling pathway. *WRKY* gene expression is downregulated by 2-deoxyglucose (2-DG) or mannose, which are inhibitors of hexokinase (HXK), and in *hxk1-3* mutants. Mutants in *WRKY17, WRKY60* or *WRKY72* have reduced hypocotyl growth in response to sucrose, but do not have altered circadian period. Our data suggest that HXK1-dependent regulation of *WRKY* genes by sugars represents a superoxide-activated transcriptional subnetwork that influences plant growth.

**Highlight:** *WRKY11, WRKY17, WRKY60* and *WRKY72* are upregulated by a sugar-activated superoxide signalling pathway in a HKX1-dependent manner. These sugar-regulated *WRKYs* represent a transcriptional subnetwork promoting plant growth.

## Introduction

Plants produce sugar from photosynthesis during the day to drive anabolism. In addition to their role as metabolic substrates and energy storage, sugars also act as signals to influence many aspects of plant physiology and development (Rolland *et al*., 2006). Photosynthesis occurs predominantly in leaf mesophyll cells, but sugar must be distributed around the plant and stored for utilisation during the night. The dependence on light to produce sugars creates specific challenges for photosynthetic organisms, particularly in natural conditions where light availability fluctuates in changeable weather or from competing neighbours (Annunziata *et al*., 2017). Thus, plants require mechanisms to sense local sugar availability and adjust metabolism and transport through signalling and regulated gene expression.

There are several well-characterised signalling pathways that contribute to sugar responses in plants. Snf1-RELATED KINASE 1 (SnRK1) is activate under carbon limitation, and its activity is inhibited by the signalling sugar trehalose-6-phosphate (T6P), which is closely correlated with concentration of sucrose (Baena-González *et al*., 2007; Zhang *et al*., 2009; Figueroa and Lunn, 2016). By contrast, TARGET OF RAPAMYCIN (TOR) kinase is activated by glucose (Xiong *et al*., 2013). Null mutations in *T6P SYNTHASE 1 (TPS1)* and essential subunits of SnRK1 or TOR are embryo lethal (Eastmond *et al*., 2002; Menand *et al*., 2002; Ramon *et al*., 2019). HEXOKINASE1 (HXK1) performs the first committed step of glycolysis, but also localises to the nucleus and has a function independent of its glycolytic activity (Moore *et al*., 2003; Cho *et al*., 2006). *HXK1* mutants are resistant to growth inhibition by high concentrations of exogenous glucose, and also grow slowly compared to wild type. The targets of both the SnRK1 and TOR kinase networks have been well described through a combination of transcriptomics, proteomics and phosphoproteomics (van Leene *et al*., 2019, 2022) whereas the signalling function of HXK1 has been less well defined.

The downstream targets of these signalling kinases include modulation of activity of transcription factors. SnRK1 can modulate basic LEUCINE ZIPPER (bZIP) transcription factors (Baena-González *et al*., 2007). For example, bZIP63 is phosphorylated by SnRK1, which alters its dimerization and DNA-binding affinity (Mair *et al*., 2015). TOR phosphorylates E2Fa transcription factor to activate cell cycle genes (Xiong *et al*., 2013) and SnRK1 phosphorylates E2Fa to promote its degradation (Son *et al*., 2022). TOR can also regulate transcription *via* histone modification by phosphorylation of FERTILIZATION-INDEPENDENT ENDOSPERM (FIE), an essential component of the POLYCOMB REPRESSOR COMPLEX 2 (PRC2) (Ye *et al*., 2022). HXK1 appears to be able to modulate both transcriptional activation and repression. In Arabidopsis, HXK1 cooperates in glucose-dependent repression by ETHYLENE INSENSITIVE 3 (EIN3) on EIN3 binding sites (Yanagisawa *et al*., 2003). In apple, HXK1 interacts with and phosphorylates bHLH3 to activate anthocyanin biosynthesis genes and anthocyanin accumulation (Hu *et al*., 2016).

Our understanding of the transcription factor networks driving responses to sugar and how they are controlled by sugar signalling is incomplete. The most prominent class of transcription factors associated with sugar responses in plants are of the bZIP family (Baena-González *et al*., 2007; Kang *et al*., 2010; Ma *et al*., 2011; Matiolli *et al*., 2011). Other examples of transcription factors include NAC (Li *et al*., 2011; Yu *et al*., 2020), Myb and Myb-related (Teng *et al*., 2005; Chen *et al*., 2017), bHLH (Stewart *et al*., 2011; Min *et al*., 2019) and WRKY (Sun *et al*., 2003; Chen *et al*., 2019; Huang *et al*., 2021).

We previously used RNA-seq in dark-adapted Arabidopsis seedlings to identify transcriptional responses to sucrose in the absence of light (Román *et al*., 2021). We found reactive oxygen species (ROS)-regulated transcripts to be a prominent feature of the response and that sugar-activated superoxide production contribute to regulation of circadian gene expression. Among the sugar-regulated genes, we detected numerous *WRKY* genes, which have commonly been associated with other ROS-associated signalling processes, such as pathogen responses and senescence (Bakshi and Oelmüller, 2014; Jiang *et al*., 2017). Here we find that these sugar-regulated *WRKY* genes act downstream of sugar-activated superoxide production and contribute to sugar-responsive hypocotyl elongation, but not regulation of the circadian clock. Regulation of these genes by sugar depends on HXK1. Our results place these WRKY transcription factors within a network of ROS-regulated sugar signalling in Arabidopsis.

## Materials and Methods

### Plant materials and growth conditions

Wild-type (Col-0), *wrky11-3* (SALK_141511), *wrky17-3* (SAIL_1230_F07), *wrky60-1* (SALK_120706) (Xu *et al*., 2006), *wrky72-2* (SALK_055293) (Bhattarai *et al*., 2010), *hxk1-3* (SALK_070739) (Lee *et al*., 2012) and *35Sp*:*LUC* (CS9966) were obtained from the Arabidopsis Biological Resource Centre. Genotypes were confirmed by PCR using the primers listed in Table S1. *DIN6p:LUC* in Col-0 has been described previously (Frank *et al*., 2018).

Seeds were surface sterilised (30% (v/v) bleach, 0.02% (v/v) Triton X-100), washed three times in sterilised water and sown on half-strength Murashige and Skoog media (1/2 MS) (Sigma), 3 mM MES-KOH (pH 5.7), solidified with 0.8% agar Type M (Sigma). Seeds were placed in the dark at 4ºC for 2 d and grown in 12 h light (∼100 µmol m^-2^ s^-1^), 12 h dark at constant 20ºC.

### *WRKY* promoter reporter lines

An upstream region from the start codon was amplified from Col-0 gDNA for *WRKY11* (1330 bp), *WRKY17* (1656 bp), *WRKY60* (1649 bp) and *WRKY72* (1623 bp) with the primers listed in Table S1 using Phusion™ High-Fidelity DNA polymerase (Thermo Scientific). PCR products were A-tailed with Taq polymerase and cloned into pCR8/GW/TOPO (Invitrogen). *WRKY* promoters were introduced into pEarleyGate301-LUC2 (Rawat *et al*., 2009) using LR Clonase II (Invitrogen). Confirmed and sequenced constructs were transformed into *Agrobacterium tumefaciens* (C58) and introduced into Col-0 Arabidopsis using floral dip (Clough and Bent, 1999). T1 transformants were identified by resistance to 25 µg/ml phosphinothricin (PPT). T2 populations segregating 3:1 for PPT resistance were carried forward to identify homozygous T3 populations for further experiments.

### Quantitative RT-PCR

Total RNA was extracted from *ca*. 30 mg snap frozen tissue with ISOLATE II RNA Plant Kit (Meridian Bioscience). cDNA was prepared from 0.5 µg DNase-treated RNA in 10 µl reactions of Tetro cDNA synthesis kit (Meridian Bioscience) using oligo(d)T primer. 10 µl PCR reactions were performed in technical duplicate with SensiFAST SYBR No-ROX (Meridian Bioscience) with 7.5 ng cDNA and 200 nM primers (Table S1) on a CFX Opus 384 Real-time PCR System (BioRad). Mean PCR reaction efficiencies were calculated for each primer pair with LinRegPCR (Ruijter *et al*., 2009) and used to calculate gene expression levels (PCR_efficiency^-Ct^). *UBQ10* was chosen as the reference gene because it was stable in previous RNA-seq experiments in equivalent conditions (Román *et al*., 2021).

### Superoxide detection

Seedlings were collected under dim green light into freshly prepared staining solution (2 mg/ml (w/v) nitroblue tetrazolium, 10 mM potassium phosphate buffer (pH 7.8), 10 mM NaN_3_) and vacuum infiltrated in the dark for 1 min. Samples were cleared by boiling for 5 min in 1:1:4 lactic acid:glycerol:ethanol then transferred to 1:4 glycerol:ethanol. Shoots were mounted on coverslips and imaged on an Epson V370 Photo flatbed scanner and stain intensity was quantied with ImageJ (NIH).

### Luciferase experiments

To measure the effect of sugars on promoter-luciferase reporter activity, pairs of 7 d old seedlings were transferred to 96-well LUMITRAC™ 200 plates (Greiner) containing 250 µl ½ MS (0.8% agar) per well and seedlings were grown in the dark for 72 h. Seedlings were treated with 1mM D-luciferin K^+^ salt (Cayman Chemicals) >12 h before measurements. At subjective dawn in dim green light, 25 µl of sugars were added and luminescence was monitored using orbital scan mode in a LUMIstar Omega plate reader (BMG Labtech).

To measure circadian rhythms in *wrky* mutants, we used a previously described Arabidopsis seedling transformation protocol (Ting *et al*., 2022) with some modifications. *A. tumefaciens* (C58) carrying *GIp:LUC2* was cultured overnight in LB, 10 mM MES-KOH (pH 5.5). Pelleted cells were resuspended in infiltration buffer (10 mM MgCl_2_, 10 mM MES-KOH pH 5.5, 200 µM acetosyringone). Five d old seedlings were vacuum infiltrated with bacterial culture (2 × 1 min) then returned to the growth cabinet. Seven d old seedlings were transferred to ½ MS media containing 50 µg/ml timentin and 1 mM D-luciferin K^+^ salt was applied. Luciferase luminescence was imaged from the following morning in continuous light (40 µmol m^-2^ red and blue LED; LB3, Photek) using a Retiga LUMO CCD camera (Teledyne Photometrics). Circadian rhythms were analysed using FFT-NLLS in Biodare (Zielinski *et al*., 2014).

## Results

We previously used RNA-seq to identify Arabidopsis genes that are regulated by sugar independently of light (Román *et al*., 2021). Briefly, seedlings were transferred to the dark for 72 h, then either treated with sucrose or mannitol in the dark or transferred to the light with or without an inhibitor of photosynthesis. Comparison of this list of sugar-regulated genes with published data of SnRK1- and TOR-regulated genes identified a list of ca. 1000 genes that were specific to our dataset and were enriched for genes associated with ROS signalling (Román *et al*., 2021). To gain insight of this sugar-regulated transcriptional network, we used this gene list to identify any over-represented family of transcription factor. Among the ten largest transcription factor families in the Arabidopsis genome (Jin *et al*., 2017), only WRKY genes were significantly enriched in this dataset (Fig 1A).

**Figure 1.**
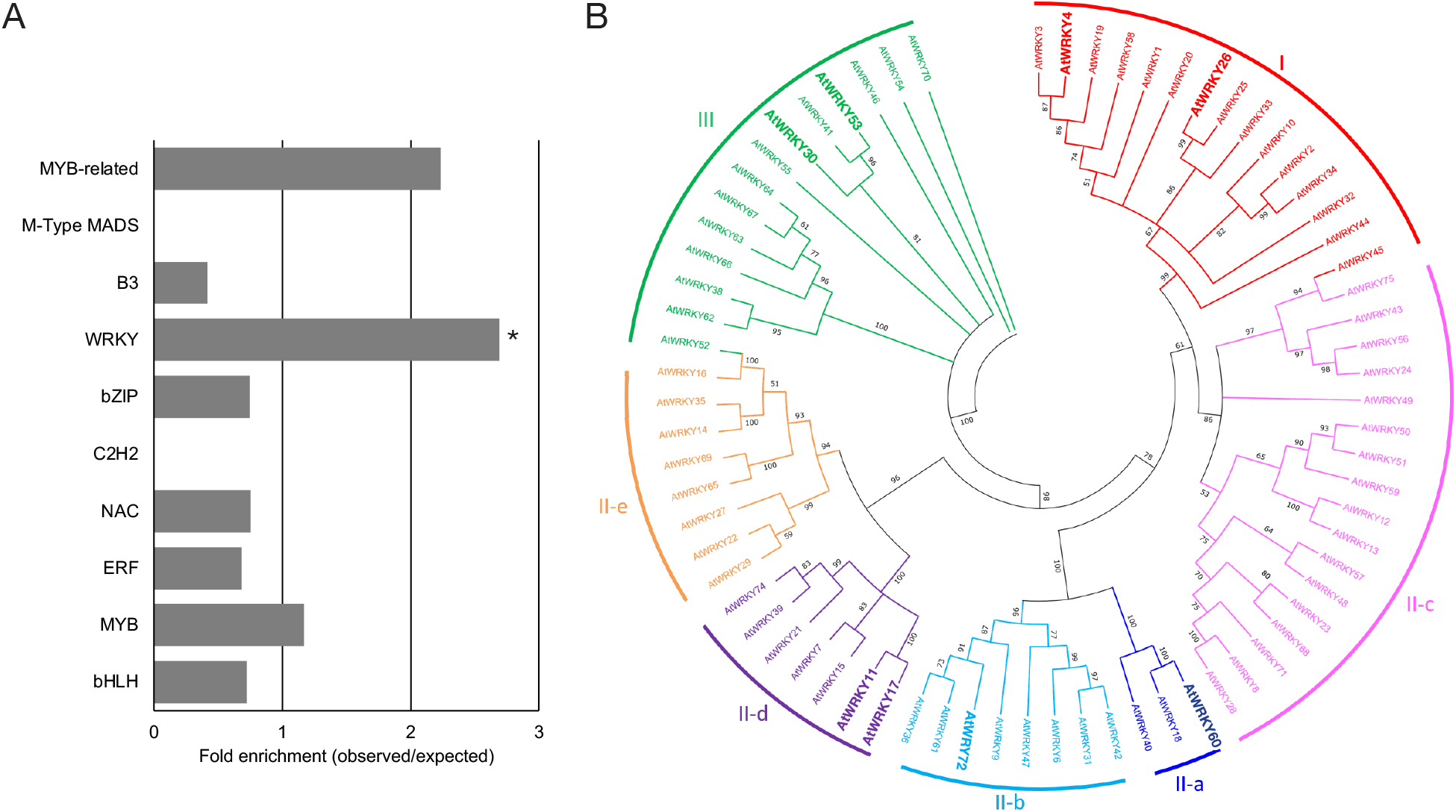
Sugar-regulated WRKY transcription factors. (A) Enrichment of genes from the ten largest transcription factor families among a published list of *ca*. 1000 SnRK1-/TOR-independent sugar-regulated genes (Román *et al*., 2021). * *P* < 0.05, χ^2^. (B) Phylogenetic tree of 72 WRKY proteins from Arabidopsis. The tree was built with IQ-Tree (v 1.6.10) using full length protein sequences (10,000 Ultrafast Bootstrap replicates, 1,000 maximum iterations, cut-off 50%) and visualised with MEGA-X (v. 10.0.5). Eight sugar-regulated WRKYs are indicated by large text.

There are 72 WRKY genes in the Arabidopsis genome and eight sugar-regulated WRKY genes were identified in the RNA-seq dataset (Fig S1). Phylogenetic analysis of the WRKY proteins shows that the eight sugar-regulated WRKYs are dispersed across five of the seven subclasses of the WRKY family (Fig 1B). *WRKY11* and *WRKY17* and the most closely related among the eight sugar-regulated *WRKY* genes and have been reported to have a functionally redundant role in pathogen responses (Somssich *et al*., 2006).

We focussed on *WRKY11, WRKY17, WRKY60* and *WRKY72* because these four genes were most strongly upregulated by sucrose in dark-adapted seedlings (Fig S1) and T-DNA mutants are available (Fig S2). Using qRT-PCR, we confirmed that these four genes were upregulated in response to sucrose, compared to mannitol controls in dark-adapted seedlings (Fig 2). Furthermore, we tested whether the induction of these genes by sucrose was inhibited by DPI or PI, two chemicals that inhibit a sugar-activated superoxide-Ca^2+^ signalling pathway (Román *et al*., 2021; Li *et al*., 2022). Upregulation of all four *WRKY* genes by sugar was significantly inhibited by either DPI or PI (Fig 2A). We used nitroblue tetrazolium (NBT) stains to test whether sucrose-induced superoxide accumulation was affected in the *wrky* mutants. Superoxide levels were significantly higher in sucrose treated seedlings in all genotypes, similar to wild type (Fig 2B,C). Together, these data suggest that these *WRKY* genes are downstream targets of this superoxide-activated signalling pathway.

**Figure 2.**
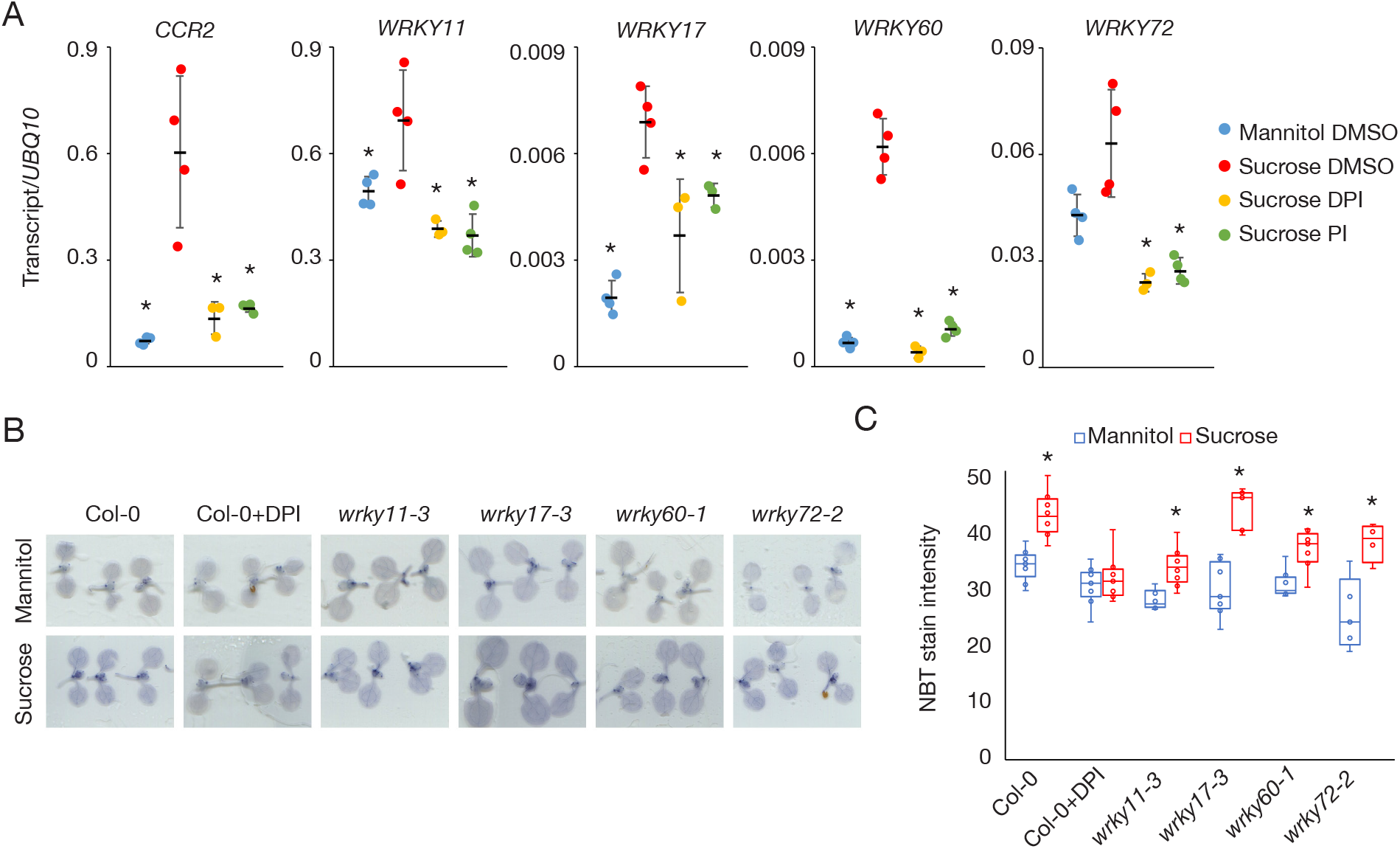
*WRKY* gene expression is downstream of sugar-activated superoxide. (A) Transcript level of *CCR2, WRKY11, WRKY17, WRKY60* and *WRKY72* relative to *UBQ10* in 10 d old Col-0 seedlings 12 h after treatment with 30 mM mannitol, 30 mM sucrose or 30 mM sucrose in the presence of 10 µM DPI or 25 µM PI at subjective dawn following 72 h in the dark (means ± SD, n = 4; * *P* < 0.05 from sucrose, Bonferroni-corrected *t*-test. (B) Images and (C) quantification of NBT stains of 10 d old wild-type Col-0, *wrky11-3, wrky17-3, wrky60-1* and *wrky72-2* seedlings treated with 30 mM mannitol or 30 mM sucrose after 3 days in the dark (Tukey’s boxplots, n = 8; * *P* < 0.05 from wild type, Bonferroni corrected *t*-test).

We have proposed that sugar-activated superoxide signalling contributes to growth because DPI and PI both inhibit hypocotyl elongation in dark-grown seedlings (Román *et al*., 2021; Li *et al*., 2022). We tested the effect of sucrose on hypocotyl growth in the *wrky* mutants. Similar to the effect of DPI and PI, the effect of sucrose on hypocotyl elongation was significantly less in *wrky17-3, wrky60-1* and *wrky72-2* mutants compared to wild type (Fig 3).

**Figure 3.**
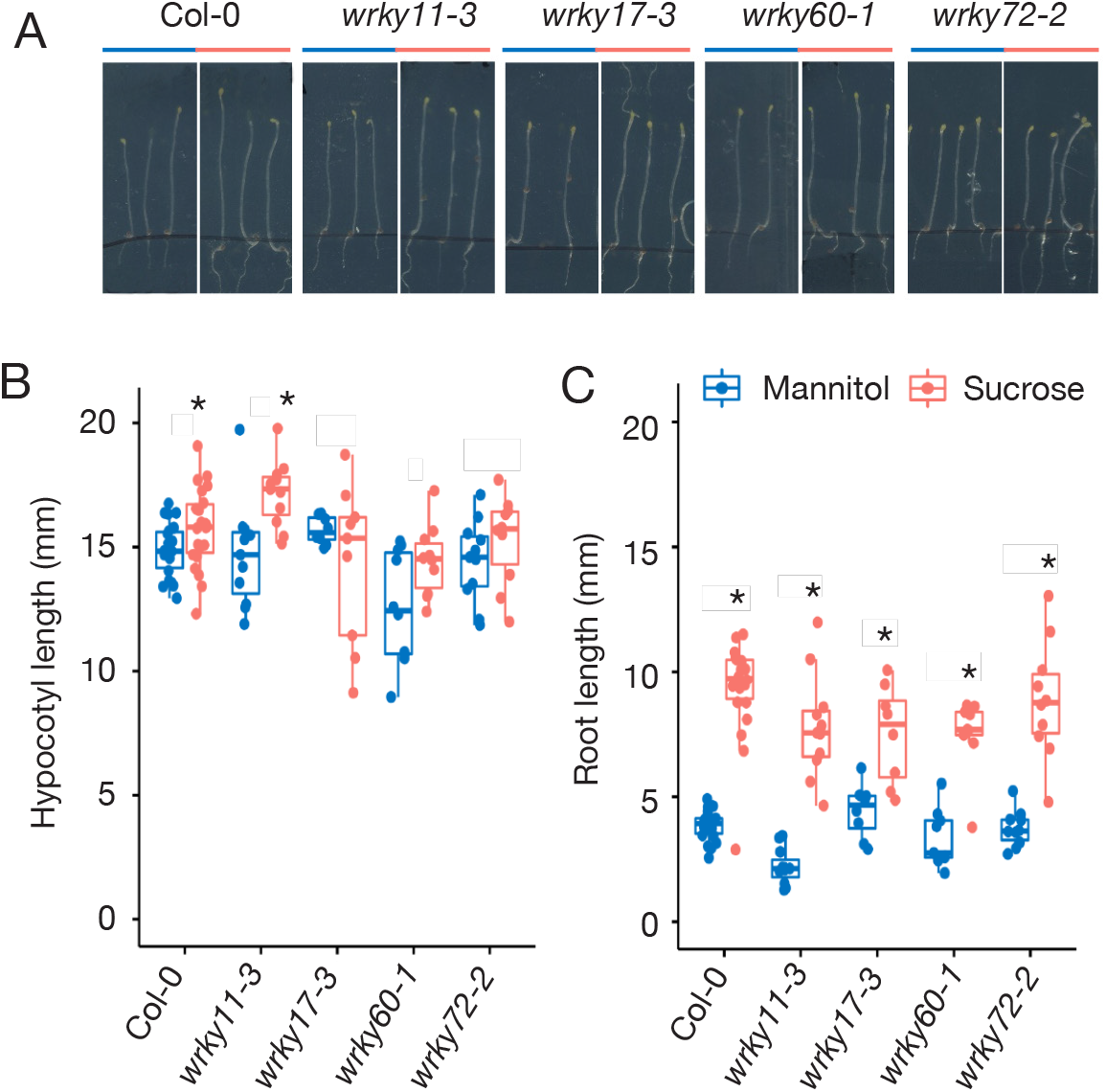
Sugar-regulated *WRKYs* are required for hypocotyl response to sucrose. (A) Images (B) hypocotyl length and (C) root length of 7 d old Col-0, *wrky11-3, wrky17-3, wrky60-1*, and *wrky72-2* grown in the dark on ½ MS with 30 mM mannitol (blue) or sucrose (red) for 5 d.

Sugar-activated superoxide influences the expression of circadian clock genes in the evening (Román *et al*., 2021; Chen *et al*., 2022). We therefore considered whether sugar-regulated *WRKY* genes are part of the circadian system in Arabidopsis. We used a published circadian transcriptome to identify whether any *WRKY* genes have detectable rhythms of expression in continuous light (Romanowski *et al*., 2020). We identified a total of six circadian-regulated *WRKY* genes (Fig 4A). Among them were *WRKY4* and *WRKY26*, which were downregulated by sugar in our RNA-seq experiment (Fig S1), *WRKY18* which has been reported to contribute to activation of a glucose-regulated genes (Chen *et al*., 2019), and *WRKY11* and *WRKY17*. Significant rhythms were not detected for *WRKY60* or *WRKY72*.

**Figure 4.**
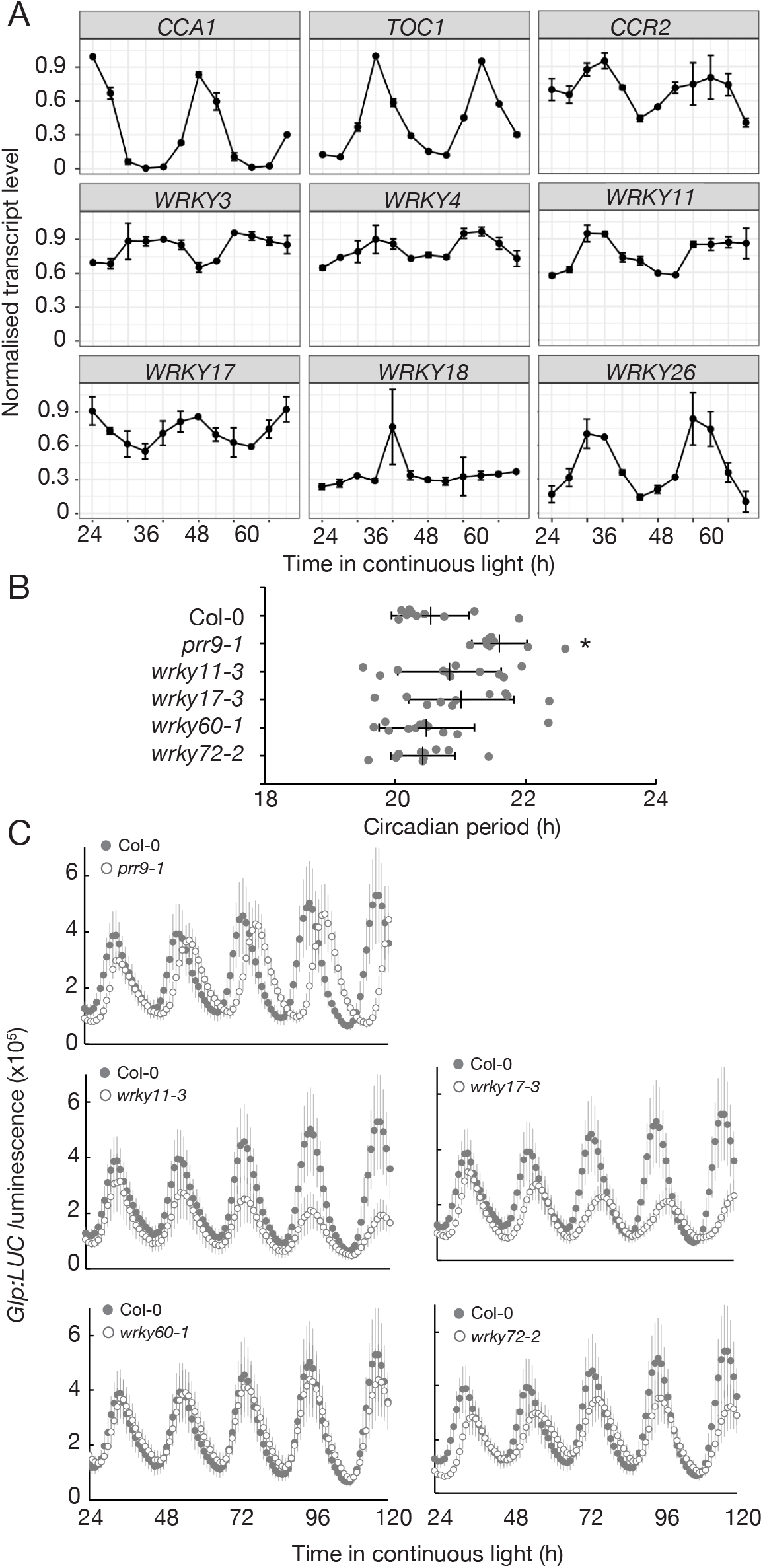
Sugar-regulated *WRKYs* do not influence circadian period. (A) Transcript level of six *WRKY* genes identified as significantly rhythmic in continuous light (Romanowski *et al*., 2020). Circadian genes *CCA1, TOC1* and *CCR2* are shown for comparison (means ± SD, n = 2). (B) Luciferase luminescence and (C) period estimates of *GIp:LUC* in wild type Col-0, *prr9-1, wrky11-3, wrky17-3, wrky60-1* and *wrky72-2* (means ± SD, n = 8; *P* < 0.05, Bonferroni-corrected *t*-test).

Since there are circadian rhythms of some sugar-regulated *WRKY* genes, and DPI and PI both lengthen circadian period (Li *et al*., 2022), we considered whether the *wrky* mutants had altered circadian period. We introduced a *GIp:LUC* reporter into Arabidopsis seedlings using Agrobacterium-mediated infiltration (Ting *et al*., 2022) and measured rhythms of luciferase luminescence in continuous light (Fig 4B). We did not detect a significant difference in circadian period between any of the four *wrky* mutants compared to wild type. This suggests that although there are modest circadian rhythms of *WRKY11* and *WRKY17* gene expression, sugar-regulated *WRKYs* do not influence circadian period.

In order to more closely examine the regulation of *WRKY* genes by sugar, we generated promoter:LUC reporters for *WRKY11, WRKY17, WRKY60* and *WRKY72* and generated stable transgenic Arabidopsis lines. We confirmed that sucrose increased luciferase reporter activity in multiple, independent transgenic for all four reporters, as expected (Fig 5A). The increase in reporter activity by sucrose compared to mannitol was detectable within several hours of sugar application, reaching a peak within about six hours for *WRKY11, WRKY17* and *WRKY60* and after about 12 hours for *WRKY72. WRKY11p:LUC* was the least responsive and *WRKY60p:LUC* was the most responsive to sucrose, consistent with the qRT-PCR results (Fig 2A). This suggests that the reporters are accurately reporting promoter activity for these *WRKY* genes.

**Figure 5.**
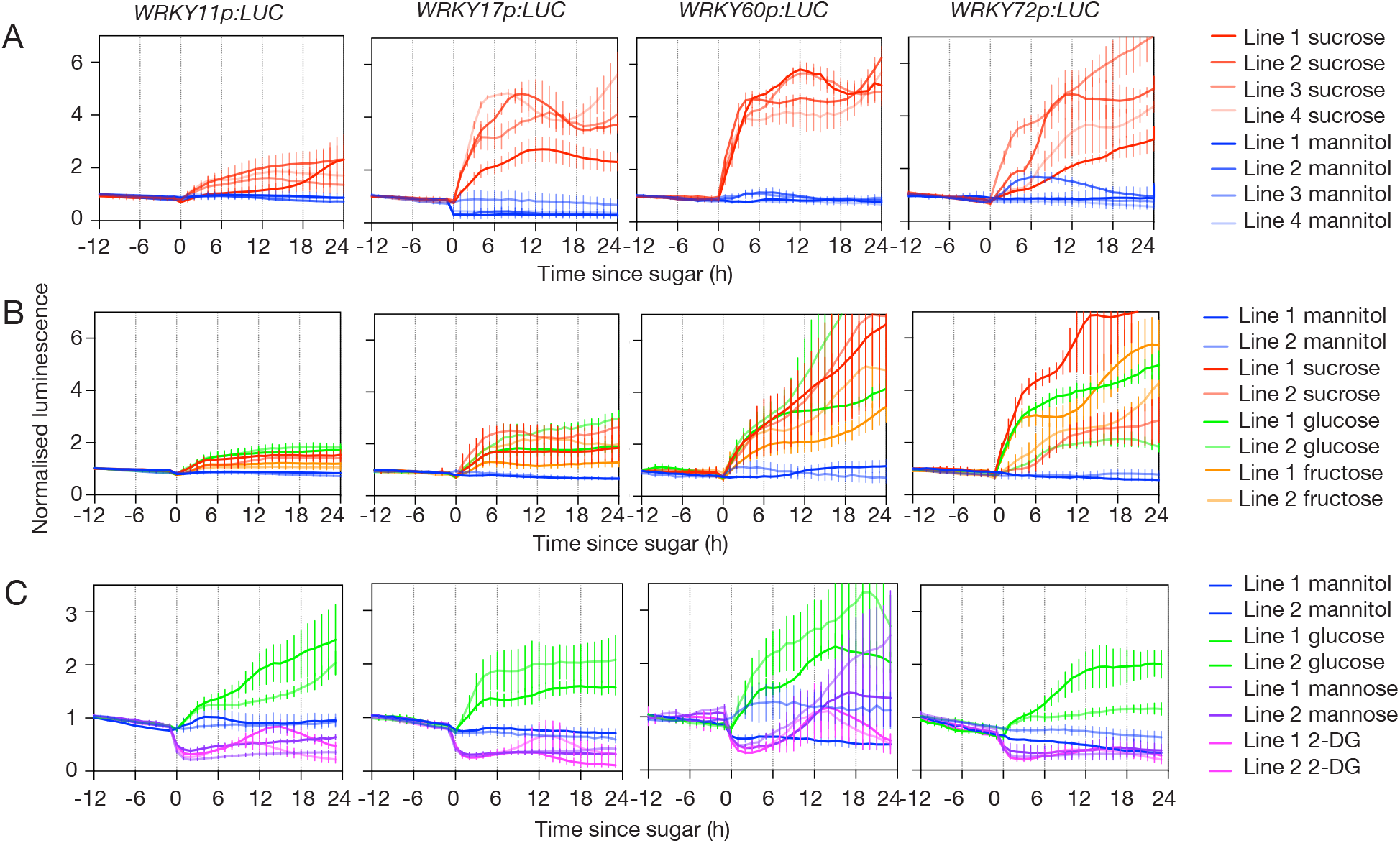
Hexokinase inhibitors reduce *WRKY* promoter activity. Normalised luminescence in dark-adapted transgenic Col-0 seedlings with *WRKYp:LUC* reporters treated at subjective dawn with (A) 30 mM sucrose or mannitol, (B) 30 mM mannitol, glucose, fructose or sucrose or (C) 30 mM mannitol, glucose, mannose or 2-DG (means ± SD, n = 4).

We next used the *WRKY* reporter lines to test the effect of sugars other than sucrose. The effect of glucose and fructose were similar to sucrose (Fig 5B), suggesting *WRKY* gene regulation is not sucrose-specific. Non-metabolisable sugars sorbitol, 3-O-methylglucose (3-OMG) and sucralose did not influence *WRKY* report activity, similar to mannitol (Fig S3). However, mannose and 2-deoxyglucose (2-DG) both rapidly reduced activity of all four reporter lines (Fig 5C). We confirmed this effect was not due to inhibition of luciferase activity using 35Sp:LUC seedlings (Fig S4).

Mannose and 2-DG can be phosphorylated by HXK and inhibit activity (Pego *et al*., 1999). Therefore, the inhibition of *WRKY* reporter activity by these sugars might suggest that HXK1 contributes to activation of *WRKY* gene expression by sucrose. We used the *WRKY* reporter lines to test the effect of 2-DG and mannose on activation by sucrose and detected lower luciferase activity for all four reporters in the presence of either inhibitor (Fig 6A). To corroborate this result, we used qRT-PCR to measure levels of *WRKY* transcripts in *hxk1-3* mutants (Fig 6B). The increase in *WRKY* transcripts in dark-adapted seedlings treated with sucrose compared to mannitol was significantly reduced in *hxk1-3* compared to wild type. This is in contrast to the circadian-regulated gene, *CCR2*, which is activated similarly in wild type and *hxk1-3*. This is consistent with previous results (Li *et al*., 2022) and suggests that HXK1-dependent *WRKY* gene regulation is distinct from HXK1-independent circadian gene regulation.

**Figure 6.**
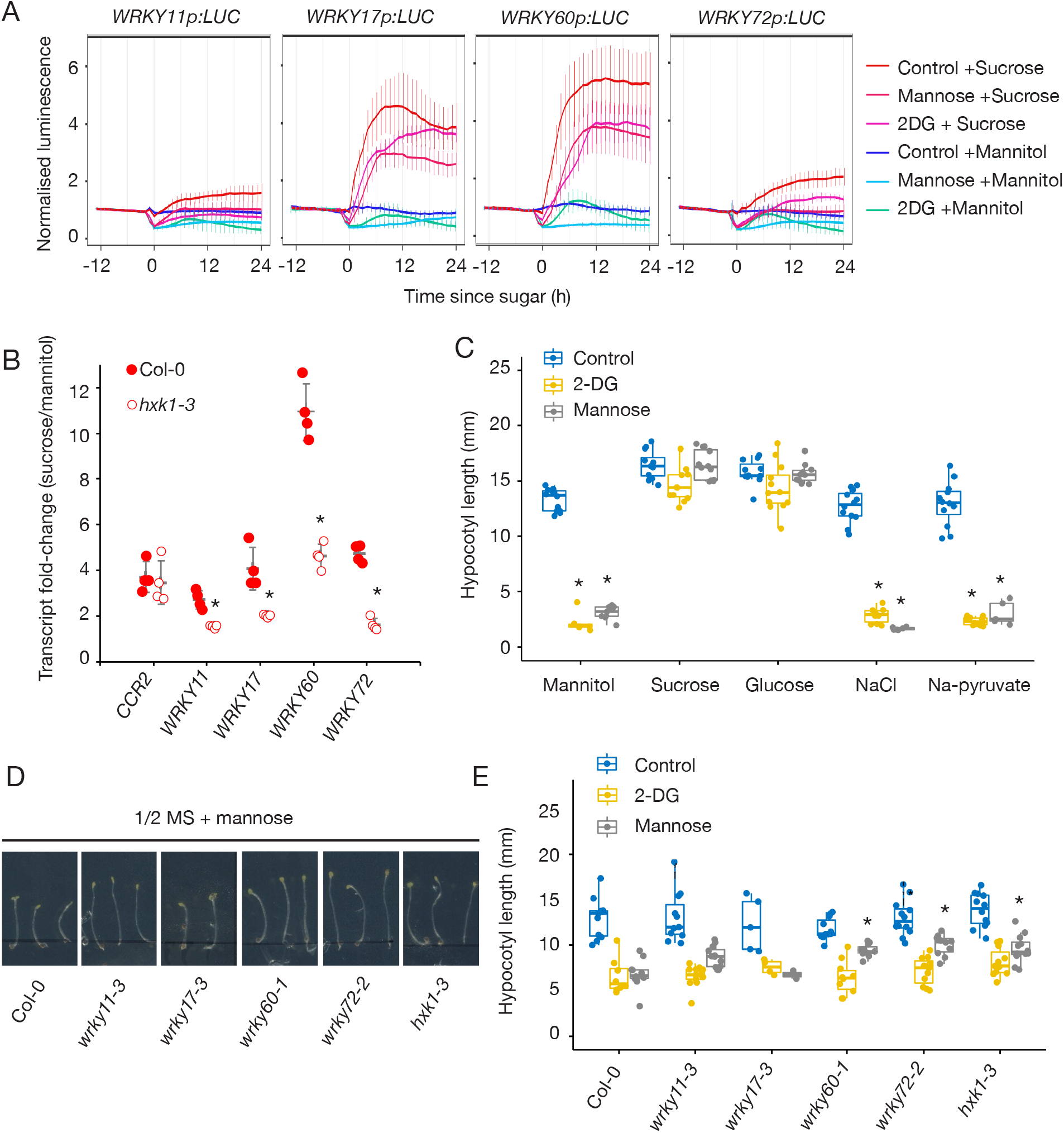
HXK1 contributes to upregulate *WRKY* genes by sugar. (A) Normalised luminescence in dark-adapted *WRKYp:LUC* transgenic seedlings treated with 30 mM sucrose or mannitol in the presence of 15 mM mannose or 2-DG (means ± SD, n = 6). (B) Fold-change of *CCR2* and *WRKY* transcript levels relative to *UBQ10* in dark-adapted wild type Col-0 and *hxk1-3* seedlings treated with 30 mM sucrose compared to mannitol for 8 h (means ± SD, n =4; * *P* < 0.05 from wild type, Bonferroni-corrected *t*-test). (C) Hypocotyl length of 7 d old seedlings grown in the dark for 5 d on 30 mM mannitol, sucrose, glucose, NaCl or Na-pyruvate on control media (1/2 MS) or with 5 mM mannose or 0.5 mM 2-DG (Tukey boxplots, n = 10; * *P* < 0.05 from control, Bonferroni-corrected *t*-test). (D) Images and (E) hypocotyl length of 7 d old wild-type, *hxk1-3* and *wrky* mutant seedlings grown in the dark for 5 d on 1 mM mannose or 0.1 mM 2-DG (Tukey’s box plots, n = 8; * *P* < 0.05 from wild-type, Bonferroni-corrected *t-*test).

To test whether HXK1 influences the contribution of *WRKY* genes to hypocotyl growth, we tested the effect of mannose and 2-DG. Both chemicals were very effective at inhibiting hypocotyl elongation (Fig 6C). Surprisingly, the effect was suppressed in the presence of sucrose or glucose, but not pyruvate. Similarly, pyruvate could not suppress the effect of mannose or 2-DG on *WRKYp:LUC* reporters (Fig S5). These suggests the effect of 2DG and mannose on hypocotyl elongation and *WRKY* gene expression is not by inhibition of glycolysis, but perhaps depends on signalling function of HXK1. Furthermore, all *wrky60-1* and *wrky72-2* mutants were partially resistant to inhibition of hypocotyl growth by mannose, similar to *hxk1-3* (Fig 6D). These data suggest sugar-regulated *WRKY* genes participate in a HXK1-dependent superoxide signalling pathway influencing plant growth.

## Discussion

We have identified at least four sugar-regulated *WRKY* transcription factor genes that act downstream of a recently identified superoxide-Ca^2+^ signalling pathway. Using promoter reporters for *WRKY11, WRKY17, WRKY60* and *WRKY72* we find that they are upregulated by sucrose, glucose or fructose but downregulated by inhibitors of hexokinase. *HXK1* contributes to upregulation of all four *WRKY* genes by sugar. Promotion of hypocotyl elongation by sucrose is reduced in mutants of these *WRKY* genes, but sucrose-stimulated superoxide accumulation and circadian period are unaffected. Thus, WRKY transcription factors contribute to a specific aspect of the sugar-regulated transcriptional network triggered by this metabolic signalling pathway, which is required for sucrose to stimulate hypocotyl growth.

Sugar promotes accumulation of superoxide and cytosolic Ca^2+^ in both Arabidopsis and rice and acts to increase expression of circadian clock genes in the evening (Román *et al*., 2021; Chen *et al*., 2022; Li *et al*., 2022). Pharmacological inhibition of this superoxide-Ca^2+^ signalling pathway in Arabidopsis lengthens circadian period (Li *et al*., 2022). We found that the same inhibitors also reduce the response the *WRKY* transcripts to sucrose (Fig 2), suggesting they are also activated by the superoxide-Ca^2+^ pathway. However, we did not detect lengthened circadian period in *wrky* mutants (Fig 4). Furthermore, although upregulation of *WRKY* genes by sucrose was lower in *hxk1-3* mutants, upregulation of *CCR2*, a circadian-regulated marker gene, was not affected (Fig 2). Thus, our data indicate that there is a HXK1-dependent transcriptional subnetwork triggered by this signalling pathway that is distinct from the mechanism acting on circadian gene expression.

Mutants in *WRKY17, WRKY60* or *WRKY72* significantly impairs the response of hypocotyl growth to sucrose (Fig 3). This suggest that these transcription factors act non-redundantly, at least with respect to their role in sugar-stimulated hypocotyl growth. This might be because WRKY proteins can form heterodimers (Xu *et al*., 2006) and loss of a single monomer could influence DNA-binding specificity. Nevertheless, *WRKY11* and *WRKY17* are closely related (Fig 1) and double mutants do have a reduced pathogen susceptibility compared to single mutants (Somssich *et al*., 2006). Thus, higher order mutants among the sugar-regulated *WRKY* genes might be expected to have a broader impact on the transcriptome and sugar-regulated processes.

HXK1 can localise to the nucleus and affect transcription factor activity (Yanagisawa *et al*., 2003; Cho *et al*., 2006; Hu *et al*., 2016). HXK1 can form a regulatory complex comprised of V-ATPASE B SUBUNIT 1 (VAB1), 26S PROTEASOME AAA-ATPASE SUBUNIT RPT5B (RPT5B) and transcription factors. This nuclear HXK1 complex binds to *cis*-elements in *CHLOROPHYLL A/B BINDING PROTEIN 2 (CAB2)* and *CAB3* to repress expression in the presence of glucose (Cho *et al*., 2006). Similarly, HXK1 promotes the glucose-dependent repression by EIN3 in protoplasts (Yanagisawa *et al*., 2003). By contrast, MdHXK1 directly interacts with and phosphorylates MdbHLH3, which stabilised this transcriptional activator to upregulate anthocyanin biosynthesis genes in the presence of glucose (Hu *et al*., 2016). We have found that hexokinase inhibitors reduce *WRKY* promoter activity (Fig 5) and sucrose-induced *WRKY* gene expression is reduced in *hxk1-3* (Fig 6). Thus, HXK1 appears to contribute to sugar-dependent upregulation of *WRKYs*, similar to its function in regulating anthocyanin biosynthesis genes in apple. However, this could occur either by assisting an activator, or inhibiting a repressor.

WRKY transcription factors are commonly associated with plant processes that are known to involve ROS signalling, notably defence responses and senescence (Bakshi and Oelmüller, 2014; Jiang *et al*., 2017). *WRKY11* and *WRKY30* were previously identified among several *WRKY* genes rapidly induced by superoxide (Scarpeci *et al*., 2008). *WRKY4, WRKY11, WRKY17, WRKY26, WRKY53, WRKY60* and *WRKY72* have been previously connected to defence responses (Somssich *et al*., 2006; Xu *et al*., 2006; Lai *et al*., 2008; Bhattarai *et al*., 2010; Kanofsky *et al*., 2017). *WRKY30* and *WRKY53* contribute to regulate senescence associated genes (Miao *et al*., 2004; Besseau *et al*., 2012). Sugars can influence both plant-pathogen interactions (Chen *et al*., 2010; Yamada *et al*., 2016) and the timing of senescence (Pourtau *et al*., 2006; Wingler *et al*., 2012), so sugar-regulated *WRKY* genes might be responding directly to changing sugar levels, or sugar-associated ROS signals during these processes. Interestingly, HXK1 positively influences leaf senescence (Dai *et al*., 1999; Pourtau *et al*., 2006), which could be mediated through activation of *WRKY* genes.

From eight sugar-regulated *WRKY* genes identified from a previous RNA-seq experiment, we have shown that at least four of those, *WRKY11, WRKY17, WRKY60* and *WRKY72*, are upregulated by a sugar-activated superoxide-Ca^2+^ signalling pathway (Li *et al*., 2022). This metabolic signalling pathway regulates evening-expressed circadian clock genes (Román *et al*., 2021). Since HXK1 contributes to the activation of *WRKY* genes, but not circadian genes, this suggests a transcriptional subnetwork that contributes to promoting hypocotyl growth. These *WRKY* genes add to the growing transcriptional network controlling sugar responses, particularly associated with ROS signalling. Further work is required to identify the specific transcription factors which interact with HXK1 and the downstream regulatory targets of the WRKYs in the network.

## Supplementary Data

Fig S1. Eight sugar-regulated *WRKY* genes identified from published RNA-Seq.

Fig S2. *WRKY* transcript levels in *wrky* mutants

Fig S3. Luciferase reporter activity in multiple, independent *WRKYp:LUC* lines.

Fig S4. Effect of mannose and 2-deoxyglucose in *35Sp:LUC* seedlings.

Fig S5. Pyruvate does not suppress effect of mannose or 2-deoxyglucose on *WRKYp:LUC* reporters.

Table S1. List of primers used in this study.

## Author contributions

MJH conceived the study; JMB, RX, YL, XL, CRB and MJH designed experiments; JMB, RX, YL and XL performed experiments; JMB, RX, YL, XL and MJH analysed data; JMB, RX, YL, CRB and MJH prepared figures; MJH wrote the manuscript; JMB, XL and CRB edited the manuscript.

## Conflicts of interest

No conflicts of interest declared.

## Funding

This research was supported by a Thomas Davies Grant from the Australian Academy of Science to MJH, the JN Peters Bequest and the University of Melbourne through Melbourne Research Scholarships to XL and CRB.

## Data availability

All data are included in the main text or as supplementary information.

## Notes

### Competing Interest Statement

The authors have declared no competing interest.

