## Supplemental Figures and Tables for "HEXOKINASE-dependent regulation of WRKY transcription factors in Arabidopsis"

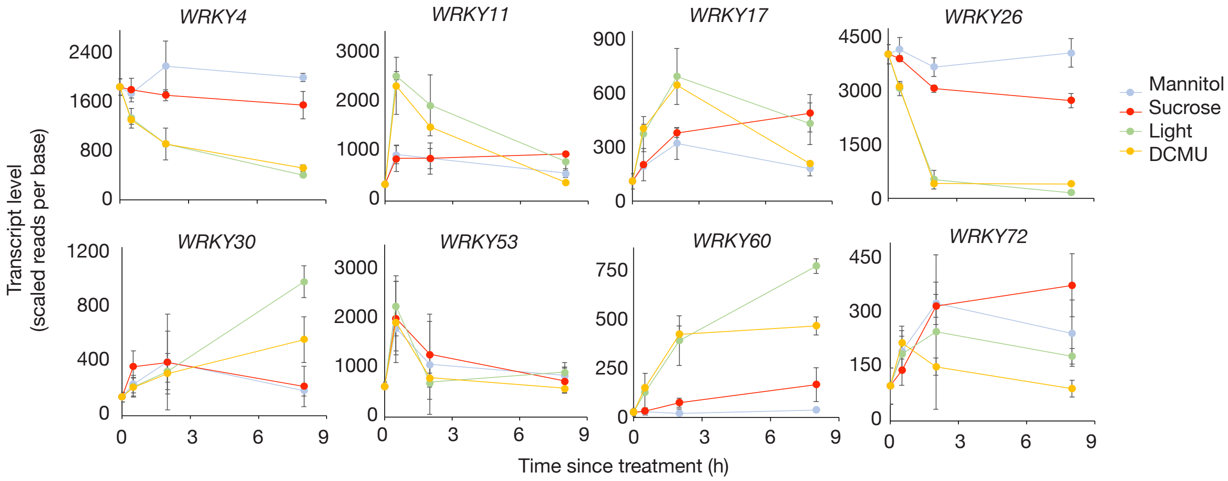


**Fig S1. Eight sugar-regulated *WRKY* genes identified from published RNA-Seq.** Transcript level of *WRKY* genes from published data (Román *et al.*, 2021). Dark adapted seedlings were transferred to 15 mM mannitol (mannitol) or sucrose (sucrose) in the dark, or transferred to the light on media with 15 mM mannitol (light) or 15 mM mannitol and 20 µM DCMU (DCMU) (means ± SD, n = 3).

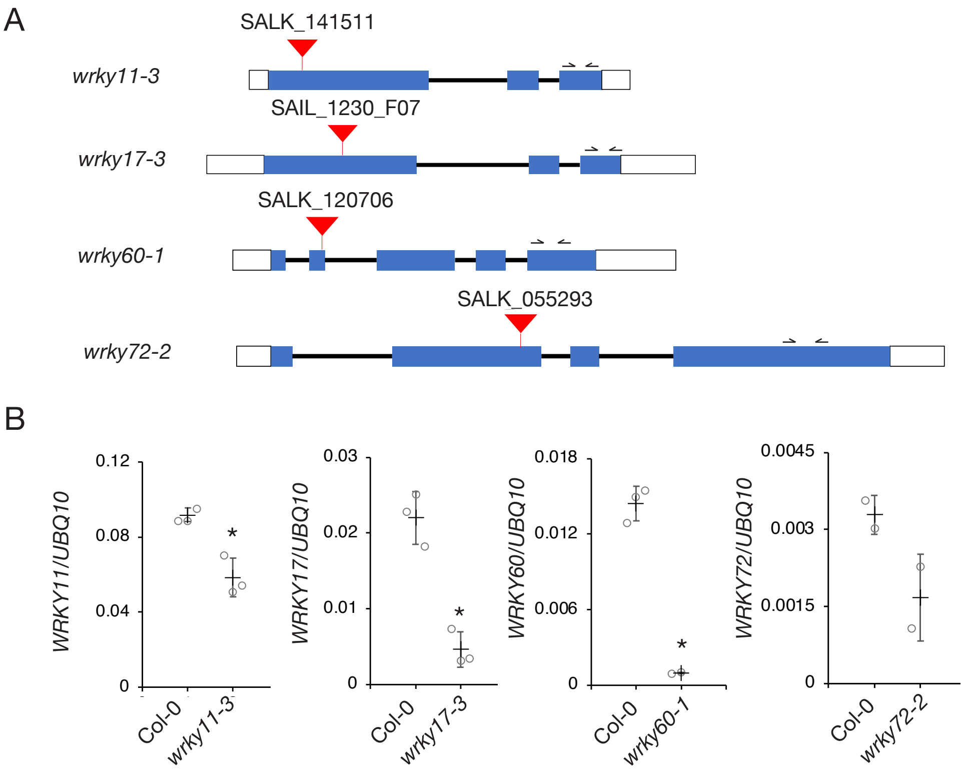


**Fig S2. *WRKY* transcript levels in *wrky* mutants.** (A) Locations of T-DNA insertions in *wrky* mutants used in this study. Protein-coding regions (blue boxes), UTRs (white boxes), introns (black lines), T-DNA insertions (red triangles) and qRT-PCR primers (arrows) are shown. (B) Transcript level of *WRKY* genes relative the *UBQ10* in wild type (Col-0) and *wrky* mutants. Means ± SD, n = 3. * $P < 0.01$, *t*-test.

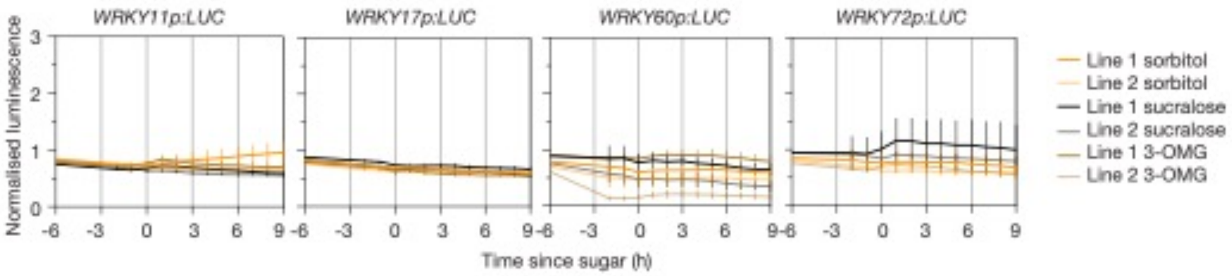


**Fig S3. Effect of non-metabolisable sugars on *WRKYp:LUC* lines.** Normalised luminescence in dark-adapted transgenic Col-0 seedlings with *WRKYp:LUC* reporters treated at subjective dawn with 30 mM sorbitol, sucralose or 3-OMG (means ± SD, n = 4).

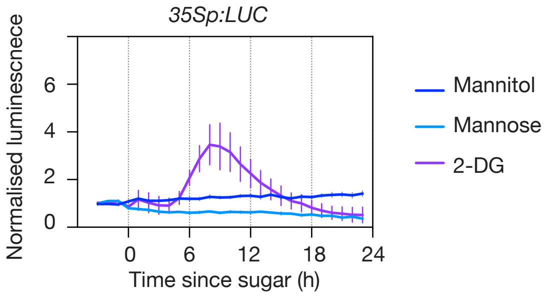


**Fig S4. Effect of mannose and 2-deoxyglucose in *35Sp:LUC* seedlings.** Normalised luminescence in dark-adapted transgenic Col-0 seedlings with a *35Sp:LUC* reporter treated at subjective dawn with 30 mM mannitol, mannose or 2-DG (means ± SD, n = 4).

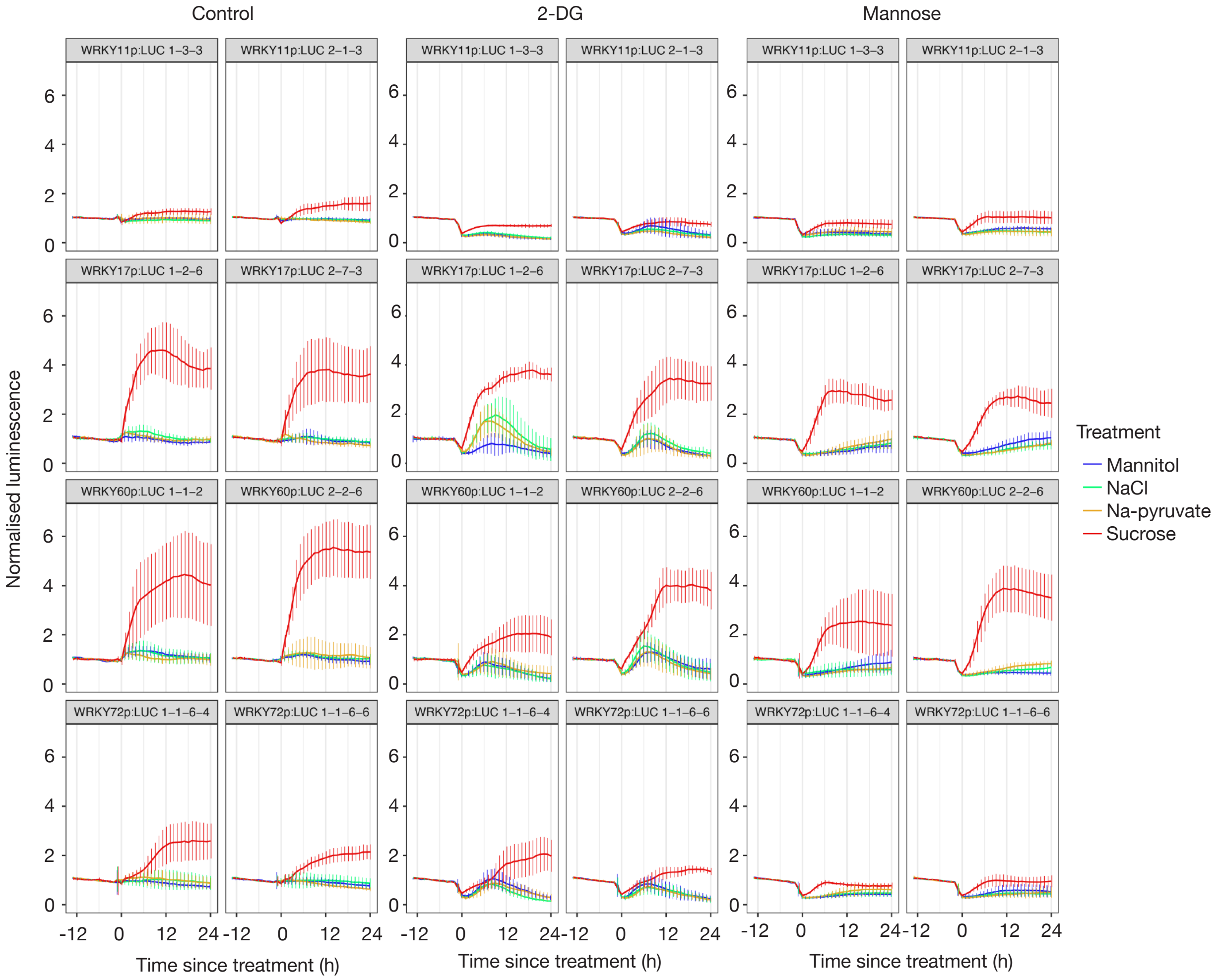


**Fig S5. Pyruvate does not suppress effect of mannose or 2-deoxyglucose on *WRKYp:LUC* reporters.** Normalised luminescence in dark-adapted transgenic Col-0 seedlings with *WRKYp:LUC* reporters treated at subjective dawn with 30 mM mannitol, sucrose, NaCl or Na-pyruvate in the presence of 15 mM mannose or 2-DG (means ± SD, n = 6).

**Table S1. List of primers used in this study.**

| Oligo name | Oligo sequence | Purpose |
|---|---|---|
| WRKY11_F | GTGGAACGAGCATTAGATGAT | qRT-PCR |
| WRKY11_R | AGCCGAGGCAAACACTAAATC | qRT-PCR |
| WRKY17_F | ACAAGTGTAGTACATTTAGAGG | qRT-PCR |
| WRKY17_R | CTAGGAGTTACATGCTCCTG | qRT-PCR |
| WRKY60_F | AGGGACACATAACCACACCG | qRT-PCR |
| WRKY60_R | TCCTCAACTGGTTCAAGCCC | qRT-PCR |
| WRKY72_F | ACAAAAGCGCTTACTTCCGA | qRT-PCR |
| WRKY72_R | TCTCCATTTGATCCGACCAT | qRT-PCR |
| CCR2_21-F | TATCGGTGCTTCGTTGGAGG | qRT-PCR |
| CCR2_74-R | CGTATTGAGCGAAGGCAGTC | qRT-PCR |
| UBQ10_F | GGCCTTGTATAATCCCTGATGAATAAG | qRT-PCR |
| UBQ10_R | AAAGAGATAACAGGAACGGAAACATAGT | qRT-PCR |
| WRKY11p_-1330F | ATGCCTTGCCTAATTTCTGTTCC | Promoter cloning |
| WRKY11p_-1R | GATGATTTCTTGGTCTGAGGATTTTG | Promoter cloning |
| WRKY17p_-1656F | ATTAGATCGAGCTGCAAATTGGC | Promoter cloning |
| WRKY17p_-1R | GATGAGAAACCAGAGGAGAAACTTG | Promoter cloning |
| WRKY60p_-1649F | ATGAAGACAAGAAAGCTGCAAGG | Promoter cloning |
| WRKY60p_-1R | AAATTTAGGTTCACAGGAGCCAA | Promoter cloning |
| WRKY72p_-1623F | AGACAGGAAAGACGACAATTTGC | Promoter cloning |
| WRKY72p_-1R | TGTCAGAAATATCAGAAGATTTTCGTC | Promoter cloning |
| wrky11-3_LP | TGTCGTATTGATGAATCGCTG | Genotyping |
| wrky11-3_RP | GTCAGTGATCTCGGAGCAGTC | Genotyping |
| wrky17-3_LP | AAAATCAGCCCAATTGTTCATAC | Genotyping |
| wrky17-3_RP | AGCAAGAAAGATCGAAGAGCC | Genotyping |
| wrky60-1_LB | CTCCAGGGCATAGTCAATGG | Genotyping |
| wrky60-1_RP | TGTTTTCGTTTCCCCGTTAG | Genotyping |
| wrky72-2_LP | GAGTGGAAGAGAGTGGCTGTG | Genotyping |
| wrky72-2_RP | CAAAACATGGTTGATCATCCC | Genotyping |
| hxk1-3_LP | TTGTTTTTGATTCCAAATCGG | Genotyping |
| hxk1-3_RP | TCATCAAATGAGGAGGAATCG | Genotyping |
| LB3 | TTCATAACCAATCTCGATACAC | Genotyping |
| LBb1.3 | ATTTTGCCGATTTCGGAAC | Genotyping |
